# The Relationship between Social Reward Behavior and Mesolimbic Dopamine Release

**DOI:** 10.1101/2025.08.11.669767

**Authors:** Travis Erickson, Henry Franks, Larry Young, Deranda Lester

## Abstract

Deficits in social behavior, such as reduced motivation and social avoidance, are key symptoms in several psychiatric disorders. Distinct modes of reward, such as drug and social, may rely on different dopamine release patterns in the mesolimbic pathway. We investigated the relationship between social reward behaviors and dopamine release elicited by phasic and tonic stimulation patterns in C57BL/6J mice. Social conditioned place preference was used to assess motivation for social interaction, and in vivo fixed potential amperometry was used to measure nucleus accumbens dopamine release before and after cocaine (10 mg/kg, ip). Additional measures included the frequency and duration of social interactions during conditioning sessions, with the first and last session representing novel and familiar social interactions respectively. No relationship was found between baseline (pre-cocaine) dopamine and social place preference in either sex. However, in males, social place preference negatively correlated with cocaine-induced phasic dopamine release, indicating that increased social motivation was associated with a reduced phasic dopaminergic response to cocaine. In contrast, greater novel interaction was associated with increased baseline dopamine elicited by tonic stimulations. These relationships were not observed in females. Overall, these findings suggest distinct, sex-dependent roles for phasic and tonic dopamine release in mediating social reward.

## Introduction

Social aversion and withdrawal are symptoms exhibited in a variety of psychiatric disorders, including depression, substance use disorder (SUD), schizophrenia, and autism (American Psychiatric Association, 2022; Yoshizumi et al., 2008). In neurotypical animals, social interaction is generally found to be rewarding (Krach et al., 2010; Venniro & Shaham, 2020), and reinforcement of social interaction has been thought to be driven by the same neural mechanisms as that of other rewarding stimuli, including drugs (Torquet et al., 2018; Hayward et al., 2018). However, emerging evidence suggests that different neural circuitry may facilitate social vs. drug reward. This study examined the relationship between neurotransmitter release classically associated with reward and motivation for social interaction.

The mesolimbic dopamine system is a crucial neurobiological circuit, typically associated with motivational functioning. The mesolimbic pathway, which includes dopamine neurons in the ventral tegmental area (VTA) that project to limbic regions including the nucleus accumbens (NAc), is thought to play a role in mediating motivation for stimuli (e.g. willingness to approach and seek stimuli) as well as reward salience (e.g. assignment of reward value on stimuli, coinciding with allocation of attention towards stimuli) (Berridge, 2007; Schultz et al., 1997). For example, the consumption of a rewarding drug increases dopamine release in the NAc, which leads to the application of incentive salience to the previously neutral cues associated with the drug and the development of motivation towards drug-seeking (Koob et al., 2016). Dysfunction of this pathway can lead to altered reward processing and maladaptive decision making, as is associated with disorders such as addiction, ADHD, anxiety/depressive disorders, and schizophrenia (Schultz et al., 1997; Serafini et al., 2020).

The mesolimbic dopamine system is a recognized part of the network of neural circuitry mediating complex social behaviors. Novel social interactions have shown to increase NAc dopamine release (Berridge & Robinson, 2003; Robinson et al., 2002; Robinson et al., 2011), and the reduction of interest in social activities is suspected to lie within a dysfunction of reward processing circuitry (Luijten et al., 2017, Volkow & Morales, 2015). However, different modes of reward have been suggested to be regulated by different accumbal-cortical circuits, potentially activated based on release properties of the stimuli-evoked dopamine (Berridge & Robinson, 2003; Bowen & Neumann, 2017). Dopamine neurons exhibit two primary patterns of activity: tonic and phasic. Tonic firing patterns (low steady firing at ∼5 Hz) set baseline dopamine levels, maintaining a constant influence on the system. In contrast, phasic firing involves brief bursts (20-100 Hz) of dopamine release, generally in response to external stimuli (Hyland et al., 2002, Dreyer et al., 2010). Phasic activity is associated with salient stimuli and has been shown to drive reward-seeking behavior more so than tonic dopamine activity (Dreyer et al., 2010; Atcherly et al., 2015). The balance between phasic and tonic activity can impact drug-related behavior. An increase in phasic firing tends to promote drug-seeking behavior, while an increase in tonic activity has the opposite effect, seemingly reducing the reinforcing effects of drugs (Budygin et al., 2020; Schultz et al., 1993).

Tonic and phasic firing patterns lead to activation of different types of dopamine receptors, classically categorized as D1 and D2. D1 receptors have a relatively low affinity for dopamine and, therefore, activation of D1 receptors often requires higher concentrations of extracellular dopamine, such as that elicited by phasic firing patterns. Activation of D1 receptors stimulates cyclic adenosine monophosphate (cAMP) signaling, which triggers a cascade of intracellular events that generally lead to neuronal excitation and spur associative learning, a crucial component in drug-seeking behavior (Caine et al., 2002; Koob & Volkow, 2016). On the other hand, D2 receptors have a high affinity for dopamine and can be activated by extracellular dopamine levels elicited from either phasic or tonic firing patterns. D2 receptor activity inhibits cAMP release, which modulates the activity of ion channels and other cellular processes, often leading to a decrease in neuronal excitability. Interestingly, D2 receptors are associated with motivation for social interaction but not deemed necessary for drug reward (Caine et al., 2002; Koob & Volkow, 2016; Liu & Wang, 2003). Instead, D2 activation has been shown to reduce drug-seeking behavior, making it a potential target for treatment (Budygin et al., 2020; Volkow & Morales, 2015). The ratio of D1 and D2 activation may be key in shifting behavior between drug and social rewards. A higher ratio of D1 to D2 activation places more emphasis on drug rewards, while a higher ratio of D2 to D1 activation attenuates the reinforcing value of drug rewards, potentially shifting focus toward social rewards (Aragona et al., 2005; Bowen & Nuemann, 2017; Koob & Volkow, 2016). Thus, reward processing hinges on the intricacies of dopamine release, including the mode of release (phasic vs tonic), as this factor dictates the activation of distinct dopamine receptor types. Drug reward is more dependent on drug-induced phasic dopamine release, while social reward is dependent on the influence of tonic dopamine release (and potentially reduced phasic dopamine release) (Bowen et al., 2017).

Motivation for social interaction has been a focus of many studies, both clinical and preclinical. Rats are a common choice of animal for preclinical studies investigating social paradigms, as rats display a greater default preference for social reward compared to other lab animals, most notably mice (Kondrakiewicz et al., 2019; Beery & Shambaugh, 2021). In addition, rats have been shown to maintain a preference for social reward even in the face of strong drug reinforcers. Kummer and colleagues (2014) compared commonly researched strains of rats (Sprague Dawley) and mice (C57BL/6) in a conditioned place preference (CPP) test designed to assess the reinforcement value of social interaction and drug (cocaine 10 mg/kg, ip). A greater percentage of rats developed social CPP relative to mice (85% of the rats vs 71% of the mice), conversely indicating that a greater percentage of the mice displayed an aversion to the social interaction relative to rats (15% of rats vs 29% of mice). When making the animals choose between stimuli associated with drug vs social reward, the relative reward strength for cocaine was 300-fold higher in mice than in rats. We used mice in this study for the translational power in the variability of mouse social behavior. We aimed to utilize the widespread social phenotypes displayed by mice (from social aversion to preference) for the purpose of better understanding the relationship between social reinforcement and aspects of mesolimbic dopamine release.

The purpose the current study was to determine the relationship between the reinforcing value of social interaction and mesolimbic dopamine functioning. We used male and female C57BL/6J mice in a social condition place preference (sCPP) task followed by *in vivo* fixed-potential amperometry for the measurement of dopamine release evoked by both phasic and tonic stimulation patterns. To assess the relationship between social preference and the dopaminergic response to a well-researched drug of abuse, mice were administered an injection of cocaine during dopamine recordings. These findings provide insight on the neurochemical mechanisms associated with drug reward salience across a social preference spectrum.

## Materials and Methods

### Animals

Sixty-one C57BL/6J mice (27 male, 34 female) were used, ranging between 3-5 months at time of experimentation. Mice were housed in same-sex groups of 2-4 and in reversed light/dark cycles with light cycle starting at 7AM and lasting 12 hours. Food and water were provided *ad libitum*.

### Social Conditioned Place Preference

Four identical plexiglass CPP apparatuses were used, each containing two removable doors that divided the apparatus into three chambers (two conditioning chambers and a neutral middle chamber). The conditioning chambers were made visually and olfactorily distinct, with different bedding materials (cardboard or aspen) and unique wall patterns (black-and-white horizontal or vertical stripes). The conditioning chambers each measured 254 mm × 254 mm each, while the middle chamber measured 101 mm × 254 mm. The current CPP procedures were adopted from those used by Kummer and colleagues (2014). Prior to entering the CPP chambers, mice habituated to the testing room in a sound-attenuation chamber for 15 min. CPP experiments occurred over 10 days, with daily 15-min sessions. All sessions were video recorded with overhead cameras. On Day 1, a preference test was conducted by placing each mouse in the neutral chamber with both doors open, allowing free movement between all three chambers. The chamber where the mouse spent the least time was designated as the non-preferred side and was paired with the social stimulus during social conditioning sessions.

Days 2, 4, 6, 8 were social conditioning days in which mice were placed in their non-preferred side along with a weight- and sex-matched novel conspecific mouse. Social behaviors that occurred during the social conditioning sessions were monitored to control for potential negative interactions between the experimental mouse and the conspecific mouse. We manually noted any fighting or aggression and used Noldus Ethovision XT video tracking software to quantify aspects of social interaction, such as the number of body contacts (frequency) and the duration of time that the mice were in close proximity of one another (within 2 cm). Novel social interactions were measured on Day 2 (the first day the experimental and conspecific mice were together), and familiar social interactions were measured on Day 8 (the fourth time the experimental and conspecific mice were together).

Days 3, 5, 7, 9 were non-social conditioning days in which mice were placed by themselves in the opposite chamber (their initially-preferred side). On Day 10, the second preference test was performed, identical to the procedures on Day 1 (Figure 1A). Social preference was determined by subtracting the time spent in the social-paired chamber on Day 10 from that of Day 1. More time spent in the social-paired chamber on Day 10 minus that of Day 1 indicates a greater preference for social interaction, suggesting a greater reinforcement value for social interaction.

**Figure 1.**
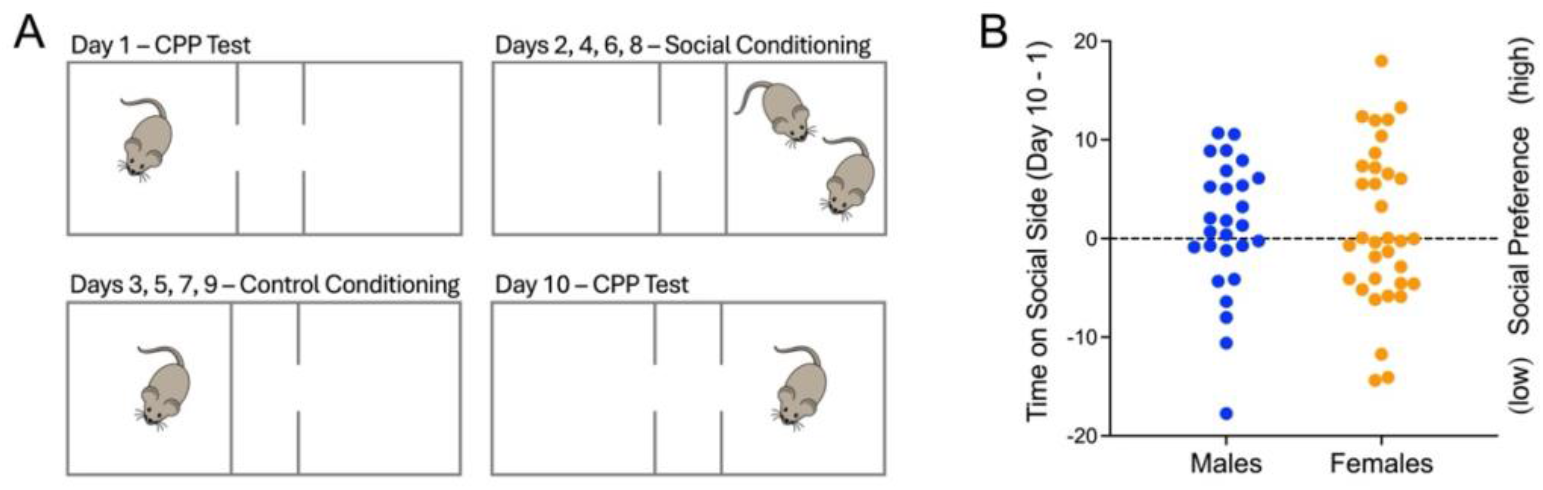
Conditioned place preference (CPP) to assess social reward. (A) The depiction of the social CPP paradigm shows the CPP test occurring first on Day 1 then again on Day 10, following conditioning days. (B) Male and female mice displayed CPP for social interaction over a similar spectrum, with mice exhibiting both high and low social preference.

### Dopamine Release Measures

Dopamine release measures took place 1-2 weeks after sCPP. Mice were anesthetized with urethane (1.5 mg/kg, i.p) prior to being placed in the stereotaxic frame (David Kompf Instruments) surrounded by a Faraday cage. The surgical procedures were similar to those of our prior studies (Estes et al., 2019; Holloway et al., 2018). Body temperature was maintained at 36 ± 0.5ºC with a heating pad. An incision was made to gain access to the skull, from which we took bregma measurements and used a mouse atlas (Paxinos and Franklin, 2001) to find proper drill sites. The bipolar stimulating electrode (SNE-100; Microprobes) was inserted into the left VTA (AP −3.3mm from bregma, ML +0.3mm from midline, and DV −4.0mm from dura), and the carbon fiber recording electrode (active recording surface of 500 µm length x 7 µm o.d.; Goodfellow) was inserted to the left NAc core (AP +1.5mm from bregma, ML +1.0mm from midline, and DV −4.0mm from dura). Lastly, the combined Ag/AgCl reference and stainless-steel auxiliary electrode was positioned contralaterally upon the cortex surface around −1.0mm from bregma.

*In vivo* fixed-potential amperometry, also known as continuous amperometry, coupled with carbon fiber recording microelectrodes has been confirmed as a valid technique for real-time monitoring of stimulation-evoked dopamine release in anesthetized rodents (Dugast et al., 1994; Forster et al., 2003; Lester et al., 2010 Holloway et al., 2018). Pharmacological studies have confirmed the measured current changes in the NAc to be dopamine-dependent (Holloway et al., 2018; Mittleman et al., 2011; Tye et al., 2013). A fixed potential of +0.8 V was applied to the brain tissue through the auxiliary electrode, and dopamine oxidation current was monitored at 10k samples per sec via the carbon fiber recording electrode and an electrometer (e-corder 401 and Picostat, eDAQ Inc) filtered at 50 Hz.

Electrical stimulations (800 µA intensity, 0.5ms pulse duration) were delivered by a programmable stimulator (Iso-Flex/Master 8; AMPI). Phasic stimulations consisted of 20 pulses at 50 Hz, and tonic simulation parameters consisted of 5 pulses at 5 Hz (Calipari et al., 2017; Fennell et al., 2020; Mahler et al., 2019). These values were based on the physiological firing properties of VTA dopamine neurons. The stimulation paradigm consisted of 5 phasic stimulations (each separated by 30 sec) then 5 tonic stimulations (each separated by 30 sec), repeated throughout the entire recording session. Following a 5 min baseline recording, the mice were injected with cocaine (10 mg/kg, ip), and amperometric recordings continued for 1 hour. Once dopamine recordings were completed, electrode sites were marked via lesioning, passing direct anodic current (100 µAmps for 10 sec) through the stimulating electrode. Mice were then euthanized (intracardial urethane 0.345 g/ml), and brains were extracted and stored for frozen cryostat sectioning and confirmation of electrode placements under a light microscope.

Carbon fiber recording electrodes were calibrated using an *in vitro* flow injection system. Dopamine standards (0.2-1.2 μM) ran across the recording electrode surface while measuring current changes. This allowed for conversion of data from current recordings (nAmp) to dopamine concentrations (uM) (Michael & Wightman, 1999; Prater et al., 2018). Dopamine release was quantified using area under the curve, and dopamine release measures post cocaine were converted into percent change from baseline (with pre-cocaine measures representing 100%).

### Data Analysis

The behavioral variables consisted of social preference (time in social chamber on Day 10 minus Day 1), frequency and duration of novel social interaction from Day 2, and frequency and duration of familiar social interaction from Day 8. The dopaminergic variables consisted of phasic and tonic baseline (pre-cocaine) dopamine release, the ratio of baseline phasic and tonic dopamine release, and phasic and tonic dopamine release 30 min post cocaine. Pearson correlations were used to assess the relationship between behavioral variables and dopaminergic variables, with alpha set at .05.

### Drugs

All chemicals were purchased from Sigma-Aldrich Chemical (St. Louis. MO). Urethane (U2500) and cocaine (C5776) was mixed in with 0.9% saline. For recording electrode calibrations, dopamine hydrochloride (H8502) was dissolved in a phosphate buffered saline (PBS, pH 7.4).

## Results

### Dopamine Release and Social Conditioned Place Preference

Social CPP was measured by subtracting the time spent in the social paired chamber on Day 10 (post conditioning) minus that of Day 1 (preconditioning). Figure 1B shows that male and female mice displayed CPP for social interaction over a similar spectrum, with mice exhibiting both high and low social preference. The wide range of social CPP scores permitted investigation of whether dopamine release correlates with individual differences in social reward preferences. Pearson correlation coefficients and corresponding p-values are listed in Table 1 for male mice and Table 2 for female mice.

**Table 1.** Males: Correlations between social behaviors and dopamine release.

|  | Social Preference | Novel Interaction Frequency | Novel Interaction Duration | Familiar Interaction Frequency | Familiar Interaction Duration |
| --- | --- | --- | --- | --- | --- |
| Phasic Dopamine Release (uM) | r = .178<br>p = .375 | r = .022<br>p = .914 | r = -.269<br>p = .174 | r = -.165<br>p = .412 | r = -.034<br>p = .868 |
| Tonic Dopamine Release (uM) | r = -.009<br>p = .966 | r = .443<br>p = .021* | r = .039<br>p = .848 | r = -.092<br>p = .647 | r = -.135<br>p = .503 |
| Phasic Release post Cocaine (%) | r = -.399<br>p = .039* | r = -.143<br>p = .476 | r = -.087<br>p = .667 | r = -.162<br>p = .418 | r = -.466<br>p = .014* |
| Tonic Release post Cocaine (%) | r = .189<br>p = .345 | r = -.563<br>p = .002* | r = -.338<br>p = .085 | r = -.219<br>p = .272 | r = -.289<br>p = .143 |

**Table 2.** Females: Correlations between social behaviors and dopamine release.

|  | Social Preference | Novel Interaction Frequency | Novel Interaction Duration | Familiar Interaction Frequency | Familiar Interaction Duration |
| --- | --- | --- | --- | --- | --- |
| Phasic Dopamine Release (uM) | r = -.079<br>p = .658 | r = -.265<br>p = .129 | r = -.157<br>p = .374 | r = .028<br>p = .875 | r = .241<br>p = .171 |
| Tonic Dopamine Release (uM) | r = .205<br>p = .245 | r = -.320<br>p = .065 | r = -.281<br>p = .107 | r = .108<br>p = .542 | r = .204<br>p = .246 |
| Phasic Release post Cocaine (%) | r = -.149<br>p = .401 | r = -.007<br>p = .970 | r = -.143<br>p = .419 | r = -.189<br>p = .285 | r = -.039<br>p = .828 |
| Tonic Release post Cocaine (%) | r = -.069<br>p = .699 | r = -.198<br>p = .263 | r = -.292<br>p = .094 | r = -.260<br>p = .138 | r = -.151<br>p = .393 |

Baseline dopamine release (prior to cocaine administration) evoked by phasic stimulations did not significantly correlate with social CPP in either males or females (Figure 2A-C). Similarly, baseline dopamine release evoked by tonic stimulations did not significantly correlate with social CPP in either males or females (Figure 2D-F).

**Figure 2.**
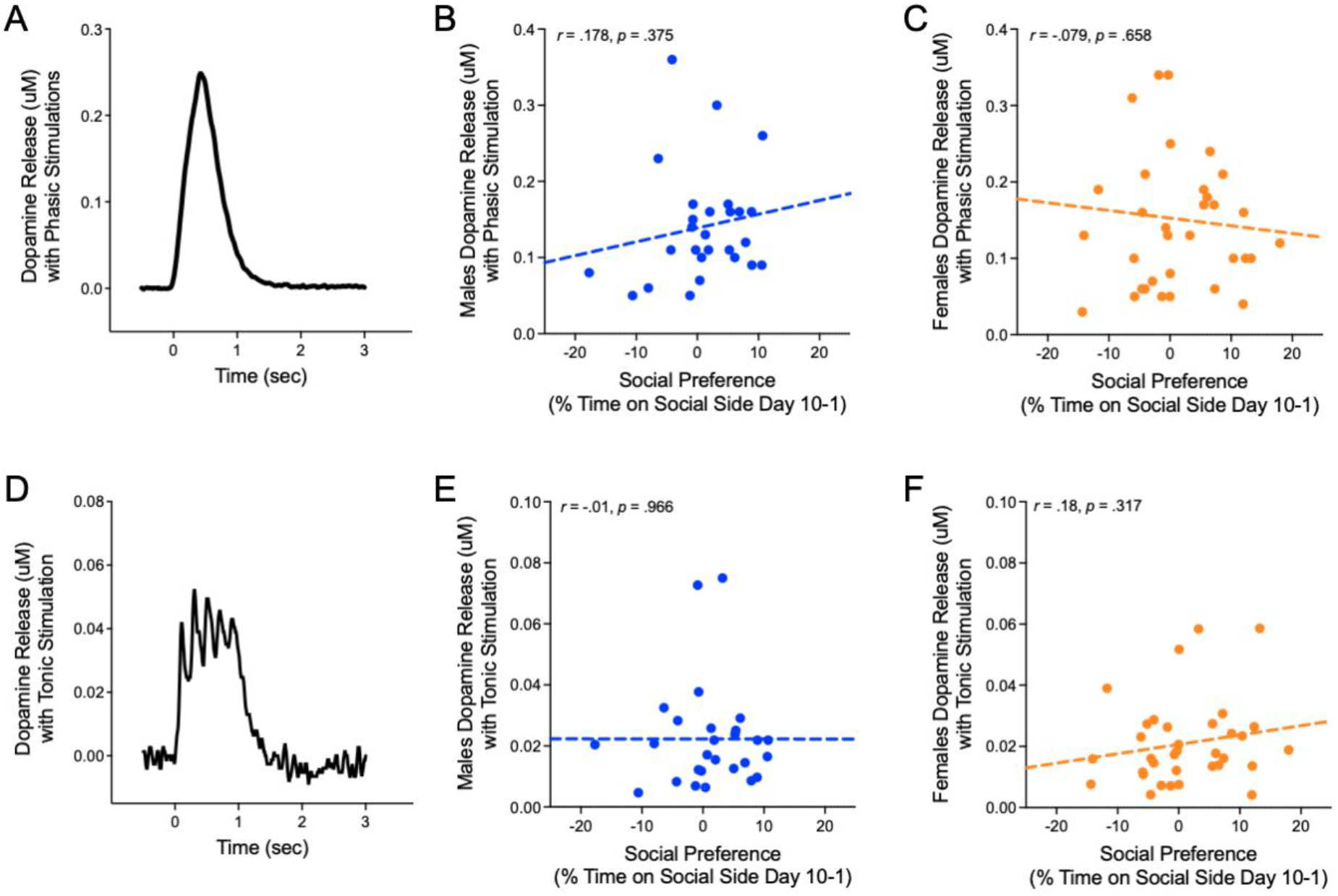
Dopamine release and correlations with social preference. (A) The example response shows dopamine release elicited by a phasic stimulation. Social preference did not significantly correlate with stimulation-evoked phasic dopamine release in (B) males or (C) females. (D) The example response shows dopamine release elicited by a tonic stimulation. Social preference did not significantly correlate with stimulation-evoked tonic dopamine release in (E) males or (F) females.

Dopamine release elicited by phasic stimulations following cocaine administration (30 min post injection) was significantly correlated with social CPP in males, but not in females (Figure 3A and B). In males, this negative relationship indicates that as social preference increases, cocaine-induced changes in phasic dopamine release decreases. Dopamine release elicited by tonic stimulations following cocaine was not significantly related to social CPP in males or females (Figure 3C and D).

**Figure 3.**
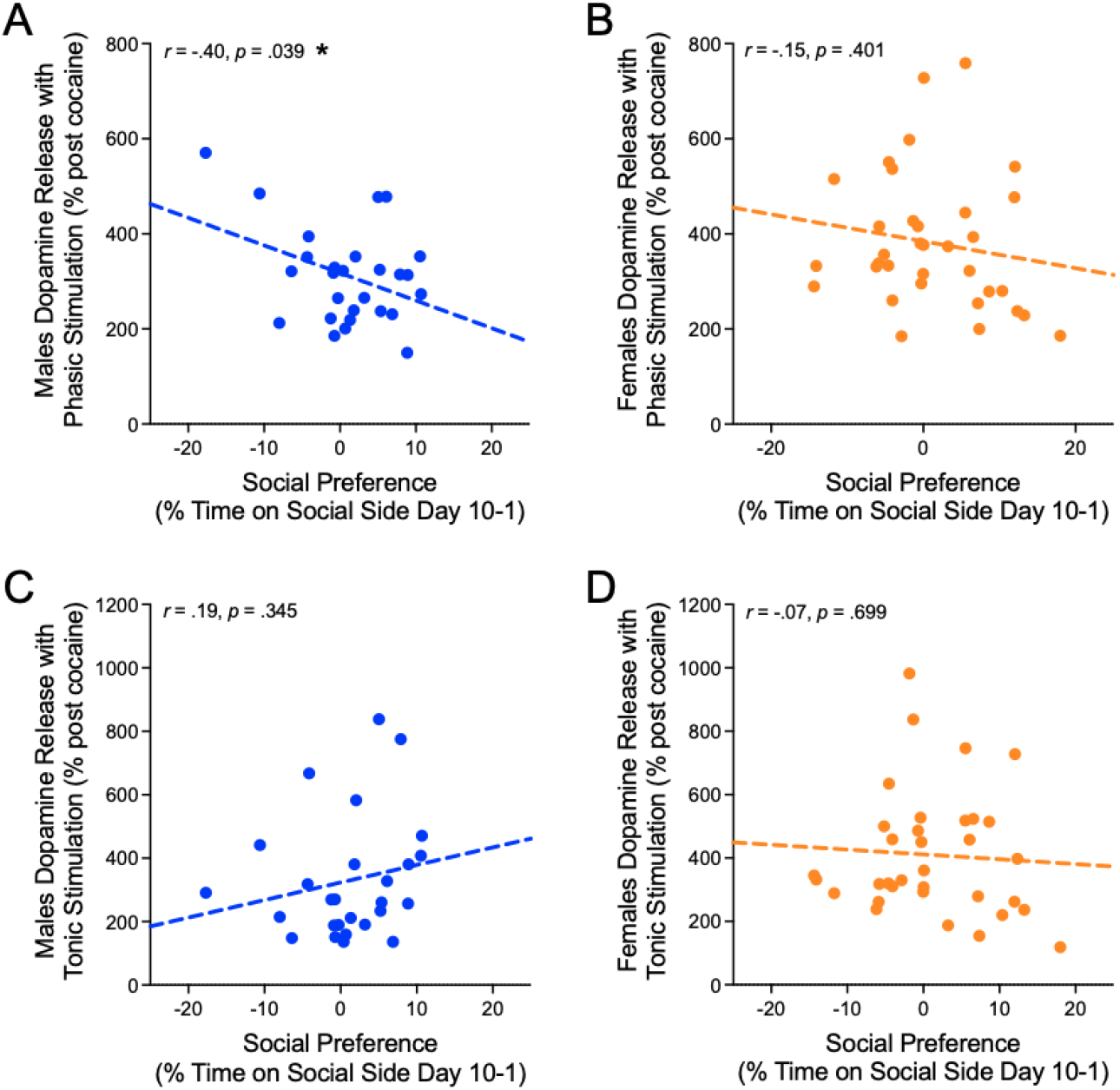
Dopamine release following cocaine correlated with social preference. We observed a significant negative relationship between social preference and phasic dopamine release post cocaine in male mice (A), but not in female mice (B). Social preference did not significantly correlate with stimulation-evoked tonic dopamine release following cocaine in (C) males or (D) females.

### Dopamine Release and Social Interaction Behaviors

Social interactions were measured by the frequency and duration of body contacts during the social conditioning session on Day 2 and Day 8, with Day 2 serving as the novel social interaction session and Day 8 the familiar social interaction session. In male mice, several significant relationships emerged between social interaction behaviors and dopamine release (Table 1, Figure 4). The frequency of novel social interactions was positively correlated with baseline tonic dopamine release (Figure 4A), indicating that males who more frequently engaged with a novel conspecific exhibited higher levels of tonic dopamine release. However, this same behavior was negatively correlated with tonic dopamine release following cocaine administration (Figure 4B), suggesting a reversal in dopamine signaling after drug exposure. In addition, the duration of familiar social interactions was negatively correlated with phasic dopamine release following cocaine (Figure 4C), indicating that males with greater engagement in familiar social interactions exhibited a blunted phasic dopamine response to cocaine. No other correlations were significant. In female mice, none of the social interaction behaviors were significantly correlated with either phasic or tonic dopamine release at baseline or following cocaine administration (Table 2).

**Figure 4.**
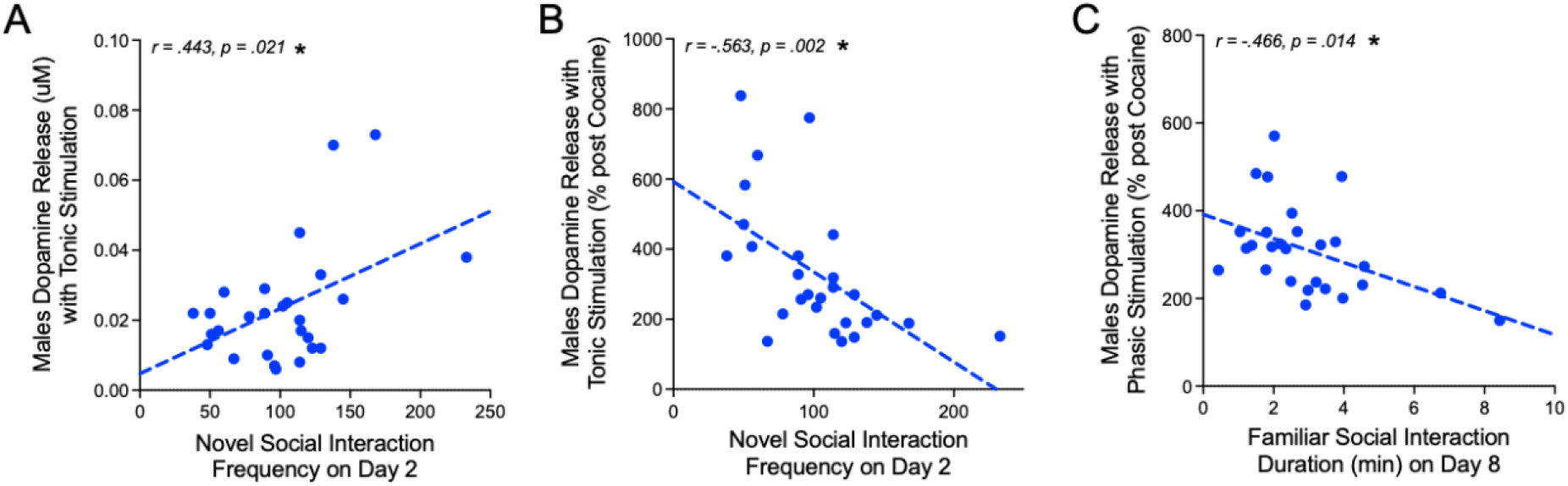
Dopamine release correlated with social interactions. In males, we observed a positive relationship between novel social interaction and tonic dopamine release (A) but a negative relationship between novel social interaction and tonic dopamine release after cocaine (B). We also observed a negative relationship between familiar social interaction and phasic dopamine release following cocaine (C).

## Discussion

A myriad of psychiatric disorders, such as substance use disorder, depression, and schizophrenia, are characterized by social deficits, including symptoms like social avoidance and withdrawal. These deficits reflect a disruption in social motivation and may involve dysfunction within the neural systems that support reward processing. The present study investigated whether naturally occurring variation in social motivation is associated with dopamine release in the NAc, a critical hub of the mesolimbic reward system, using male and female C57BL/6J mice. We correlated social behaviors (social conditioned place preference and measures of social interaction) with dopamine release before and after a cocaine injection. Dopamine release was measured using in vivo fixed-potential amperometry and electrical stimulation patterns designed to model phasic and tonic firing.

We found that the relationships between social reward behaviors and dopamine release were sex dependent. Social reward behaviors were not significantly correlated with the dopamine release measures in female mice. However, in male mice, greater social preference, as measured by social CPP, was associated with decreased phasic dopaminergic responses to cocaine, suggesting that animals with higher sensitivity to social reward show reduced dopaminergic sensitivity to drug reward. This same dopaminergic pattern was also observed in animals that spent more time interacting with a familiar conspecific, suggesting that both social preference and sustained engagement with a familiar partner are associated with blunted phasic dopamine responses to cocaine. In contrast, novel social interactions were associated with changes in tonic dopamine release. These findings add to the growing literature on the neural underpinnings of social reward and support the idea that social and drug reward processing may rely on distinct dopaminergic mechanisms.

### Social Motivation and Dopamine Release elicited by Phasic Stimulations

Among the measures of dopamine release, only phasic dopamine release following cocaine administration was significantly associated with the reinforcing value of social interaction. In male mice, social preference, defined as the change in time spent in the social chamber before and after conditioning, was significantly negatively correlated with cocaine-induced changes in phasic dopamine release. These findings suggest that in males, as social preference increases, the effects of cocaine on phasic dopamine release decreases. Inversely stated, as social preference decreases, phasic dopaminergic responses to cocaine increase. This pattern aligns with clinical findings linking social demotivation and withdrawal to increased drug-seeking and relapse risk in humans (Lookatch et al., 2019).

Phasic dopamine release plays a central role in associative learning and in assigning incentive salience to drug-predictive cues (Caine et al., 2002; Koob & Volkow, 2016); therefore, mice with lower social preference may exhibit greater cocaine-induced dopamine release, potentially reflecting increased vulnerability to drug-seeking and substance use disorder. This same dopaminergic pattern was observed in a separate measure of social motivation: the duration of interaction with a familiar conspecific. In male mice, longer interaction with a familiar partner was associated with reduced phasic dopamine release following cocaine. The convergence of these findings across both a preference-based measure (social CPP) and a direct behavioral measure (familiar interaction duration) reinforces the interpretation that strong social motivation is linked to attenuated dopaminergic responses to drug reward. While these findings are correlational, they are consistent with prior research showing that social interaction can buffer against drug-related effects and reduce vulnerability to substance use behaviors (Samperdro-Piquero et al., 2019; Venniro et al., 2018).

### Social Motivation and Dopamine Release elicited by Phasic Stimulations

Unlike phasic dopamine release, dopamine release elicited by tonic stimulation patterns was not significantly associated with social CPP, the primary measure of social reward. However, in male mice, tonic dopamine release was related to a different aspect of social behavior, the frequency of social interactions with a novel conspecific mouse. The number of social interactions during the first session (on day 2) was positively correlated with tonic dopamine release (prior to cocaine administration), indicating that greater exploratory social behavior is associated with higher tonic dopaminergic tone in males. This is consistent with previous research linking tonic dopamine levels to individual differences in novelty exploration (Frank et al., 2009). Following cocaine administration, this relationship reversed, higher novel interaction frequency was associated with lower tonic dopamine release post cocaine. This pattern mirrors the direction of the relationship observed between social preference and phasic dopamine release and may reflect a broader shift in dopaminergic responsivity to drug cues in socially motivated animals. Tonic dopamine is thought to establish the background dopaminergic tone and regulate the magnitude of phasic responses, potentially through autoreceptors mechanisms (Grace et al., 1991; Dreyer et al., 2010). Higher tonic dopamine levels have been linked to attenuated behavioral effects of cocaine and reduced drug-seeking behaviors (Budygin et al., 2020; Grace et al., 1991; Khemiri et al., 2015; Ostlund et al., 2011; Steensland et al., 2012). These findings support the idea that individual differences in social behavior relate to distinct components of mesolimbic dopamine function, and that social and drug reward may be mediated by partially opposing dopaminergic dynamics.

### Sex Differences in the Relationship Between Social Reward and Dopamine

In this study, significant associations between social behaviors and dopamine release were observed in male mice, but not in females. These sex differences suggest that social reward behaviors may be more tightly linked to dopaminergic signaling in males than in females. Clinical findings have also suggested that reward processing is more domain-specific in males compared to females (Greimel et al., 2018), and Barman et al. (2015) found that social reward anticipation and feedback engage different neural systems in a gender-specific manner, with limbic responses more strongly modulated by social traits in males. Although our study did not directly manipulate dopamine function, our results suggest that males may rely more heavily on dopamine-based mechanisms to process socially rewarding stimuli. In line with these findings, male juvenile rats have shown to be more sensitive than females to dopamine receptor antagonism during social play (Bredewold et al., 2018). Sex differences in the relationship between social reward processing and mesolimbic dopamine functioning may reflect interactions with neuromodulatory inputs to the VTA, such as oxytocin and vasopressin (Borland et al., 2018). Given our results and supporting evidence from prior studies, future research should directly examine the neurochemical mechanisms underlying sex differences in social reward processing.

## Conclusion

These findings suggest that individual differences in social behavior are linked to distinct patterns of dopamine release in the NAc, particularly in males. Specifically, greater social motivation was related to reduced phasic dopamine release in response to cocaine in male mice. Social motivation, as measured with social CPP, did not correlate with dopamine release elicited by tonic stimulation patterns; however, in males, novel social interaction was associated with increased tonic dopamine release. These results indicate that social and drug reward processing may engage overlapping but functionally distinct dopaminergic mechanisms, with potential implications for understanding protective factors against substance use. Furthermore, the observed sex differences highlight the importance of considering sex as a biological variable in studies of social reward. Future research should aim to clarify the mechanisms by which dopamine and other neuromodulators interact to support sex-specific patterns of social behavior and vulnerability to psychiatric disorders.

## References

American Psychiatric Association. (2022). Diagnostic and Statistical Manual of Mental Disorders (5th ed., text rev.). doi: 10.1176/appi.books.9780890425787

Aragona, B. J., Liu, Y., Yu, Y. J., Curtis, J. T., Detwiler, J. M., Insel, T. R., & Wang, Z. (2006). Nucleus accumbens dopamine differentially mediates the formation and maintenance of monogamous pair bonds. Nature neuroscience, 9(1), 133–139. doi: 10.1038/nn1613

Atcherley, C. W., Wood, K. M., Parent, K. L., Hashemi, P., & Heien, M. L. (2015). The coaction of tonic and phasic dopamine dynamics. Chemical Communications, 51(12), 2235–2238. doi: 10.1039/c4cc06165a

Cann, C., Venniro, M., Hope, B. T., & Ramsey, L. A. (2020). Parametric investigation of social place preference in adolescent mice. Behavioral neuroscience, 134(5), 435–443. doi: 10.1037/bne0000406

Barman, A., Richter, S., Soch, J., Deibele, A., Richter, A., Assmann, A., Wüstenberg, T., Walter, H., Seidenbecher, C. I., & Schott, B. H. (2015). Gender-specific modulation of neural mechanisms underlying social reward processing by Autism Quotient. Social Cognitive and Affective Neuroscience, 10(11), 1537–1547. doi: 10.1093/scan/nsv044

Beery, A. K., & Shambaugh, K. L. (2021). Comparative Assessment of Familiarity/Novelty Preferences in Rodents. Frontiers in Behavioral Neuroscience, 15, 648830. doi: 10.3389/fnbeh.2021.648830

Berridge, K. C., & Robinson, T. E. (2003). Parsing reward. Trends in neurosciences, 26(9), 507–513. doi: 10.1016/S0166-2236(03)00233-9

Berridge, K. C. (2007). The debate over dopamine’s role in reward: the case for incentive salience. Psychopharmacology, 191, 391–431. doi: 10.1007/s00213-006-0578-x

Borland, J. M., Grantham, K. N., Aiani, L. M., Frantz, K. J., & Albers, H. E. (2018). Role of oxytocin in the ventral tegmental area in social reinforcement. Psychoneuroendocrinology, 95, 128–137. doi: 10.1016/j.psyneuen.2018.05.028

Bowen, M. T., & Neumann, I. D. (2017). Rebalancing the Addicted Brain: Oxytocin Interference with the Neural Substrates of Addiction. Trends in Neurosciences, 40(12), 691–708. doi: 10.1016/j.tins.2017.10.003

Bredewold, R., Nascimento, N. F., Ro, G. S., Cieslewski, S. E., Reppucci, C. J., & Veenema, A. H. (2018). Involvement of dopamine, but not norepinephrine, in the sex-specific regulation of juvenile socially rewarding behavior by vasopressin. Neuropsychopharmacology, 43(10), 2109–2117. doi: 10.1038/s41386-018-0100-2

Budygin, E. A., Bass, C. E., Grinevich, V. P., Deal, A. L., Bonin, K. D., & Weiner, J. L. (2020). Opposite Consequences of Tonic and Phasic Increases in Accumbal Dopamine on Alcohol-Seeking Behavior. iScience, 23(3), 100877. doi: 10.1016/j.isci.2020.100877

Caine, S. B., Negus, S. S., Mello, N. K., Patel, S., Bristow, L., Kulagowski, J., Vallone, D., Saiardi, A., & Borrelli, E. (2002). Role of dopamine D2-like receptors in cocaine selfadministration: studies with D2 receptor mutant mice and novel D2 receptor antagonists. The Journal of Neuroscience, 22(7), 2977–2988. doi: 10.1523/JNEUROSCI.22-07-02977.2002

Calipari, E. S., Juarez, B., Morel, C., Walker, D. M., Cahill, M. E., Ribeiro, E., Roman-Ortiz, C., Ramakrishnan, C., Deisseroth, K., Han, M. H., & Nestler, E. J. (2017). Dopaminergic dynamics underlying sex-specific cocaine reward. Nature Communications, 8, 13877. doi: 10.1038/ncomms13877

Dreyer, J. K., Herrik, K. F., Berg, R. W., & Hounsgaard, J. D. (2010). Influence of phasic and tonic dopamine release on receptor activation. The Journal of Neuroscience, 30(42), 14273–14283. doi: 10.1523/JNEUROSCI.1894-10.2010

Dugast, C., Suaud-Chagny, M. F., & Gonon, F. (1994). Continuous in vivo monitoring of evoked dopamine release in the rat nucleus accumbens by amperometry. Neuroscience, 62(3), 647–654. doi: 10.1016/0306-4522(94)90466-9

Estes, M. K., Freels, T. G., Prater, W. T., & Lester, D. B. (2019). Systemic oxytocin administration alters mesolimbic dopamine release in mice. Neuroscience, 408, 226–238. doi: 10.1016/j.neuroscience.2019.04.006

Fennell, A. M., Pitts, E. G., Sexton, L. L., & Ferris, M. J. (2020). Phasic Dopamine Release Magnitude Tracks Individual Differences in Sensitization of Locomotor Response following a History of Nicotine Exposure. Scientific Reports, 10(1), 173. doi: 10.1038/s41598-019-56884-z

Forster, G. L., & Blaha, C. D. (2003). Pedunculopontine tegmental stimulation evokes striatal dopamine efflux by activation of acetylcholine and glutamate receptors in the midbrain and pons of the rat. The European journal of neuroscience, 17(4), 751–762. doi: 10.1046/j.1460-9568.2003.02511.x

Frank, M. J., Doll, B. B., Oas-Terpstra, J., & Moreno, F. (2009). Prefrontal and striatal dopaminergic genes predict individual differences in exploration and exploitation. Nature Neuroscience, 12(8), 1062–1068. doi: 10.1038/nn.2342

Franklin KB, Paxinos G. The mouse brain in stereotaxic coordinates. San Diego: Academic Press, 1997.

Grace A. A. (1991). Phasic versus tonic dopamine release and the modulation of dopamine system responsivity: a hypothesis for the etiology of schizophrenia. Neuroscience, 41(1), 1–24. doi: 10.1016/0306-4522(91)90196-u

Greimel, E., Bakos, S., Landes, I., Töllner, T., Bartling, J., Kohls, G., & Schulte-Körne, G. (2018). Sex differences in the neural underpinnings of social and monetary incentive processing during adolescence. Cognitive, Affective & Behavioral Neuroscience, 18(2), 296–312. doi: 10.3758/s13415-018-0570-z

Hayward, D. A., Pereira, E. J., Otto, A. R., & Ristic, J. (2018). Smile! Social reward drives attention. Journal of Experimental Psychology: Human Perception and Performance, 44(2), 206–214. doi: 10.1037/xhp0000459

Holloway, Z. R., Freels, T. G., Comstock, J. F., Nolen, H. G., Sable, H. J., & Lester, D. B. (2018). Comparing phasic dopamine dynamics in the striatum, nucleus accumbens, amygdala, and medial prefrontal cortex. Synapse, 73(2). doi: 10.1002/syn.22074

Hyland, B. I., Reynolds, J. N. J., Hay, J., Perk, C. G., & Miller, R. (2002). Firing modes of midbrain dopamine cells in the freely moving rat. Neuroscience, 114, 475–492. doi:10.1016/s0306-4522(02)00267-1

Khemiri, L., Steensland, P., Guterstam, J., Beck, O., Carlsson, A., Franck, J., & Jayaram-Lindström, N. (2015). The effects of the monoamine stabilizer (−)-OSU6162 on craving in alcohol dependent individuals: A human laboratory study. European neuropsychopharmacology : the journal of the European College of Neuropsychopharmacology, 25(12), 2240–2251. doi: 10.1016/j.euroneuro.2015.09.018

Koob, G. F., & Volkow, N. D. (2016). Neurobiology of addiction: a neurocircuitry analysis. The Lancet. Psychiatry, 3(8), 760–773. doi: 10.1016/S2215-0366(16)00104-8

Kondrakiewicz, K., Kostecki, M., Szadzińska, W., & Knapska, E. (2019). Ecological validity of social interaction tests in rats and mice. Genes, Brain, and Behavior, 18(1), e12525. doi: 10.1111/gbb.12525

Krach, S., Paulus, F. M., Bodden, M., & Kircher, T. (2010). The rewarding nature of social interactions. Frontiers in Behavioral Neuroscience, 4, 22. doi: 10.3389/fnbeh.2010.00022

Kummer, K. K., Hofhansel, L., Barwitz, C. M., Schardl, A., Prast, J. M., Salti, A., El Rawas, R., & Zernig, G. (2014). Differences in social interaction-vs. cocaine reward in mouse vs. rat. Frontiers in Behavioral Neuroscience, 8, 363. doi: 10.3389/fnbeh.2014.00363

Lester, D. B., Rogers, T. D., & Blaha, C. D. (2010). Acetylcholine-dopamine interactions in the pathophysiology and treatment of CNS disorders. CNS Neuroscience & Therapeutics, 16(3), 137–162. doi: 10.1111/j.1755-5949.2010.00142.x

Lookatch, S. J., Wimberly, A. S., & McKay, J. R. (2019). Effects of Social Support and 12-Step Involvement on Recovery among People in Continuing Care for Cocaine Dependence. Substance Use & Misuse, 54(13), 2144–2155. doi: 10.1080/10826084.2019.1638406

Liu, Y., & Wang, Z. X. (2003). Nucleus accumbens oxytocin and dopamine interact to regulate pair bond formation in female prairie voles. Neuroscience, 121(3), 537–544. doi: 10.1016/s0306-4522(03)00555-4

Luijten, M., Schellekens, A. F., Kühn, S., Machielse, M. W., & Sescousse, G. (2017). Disruption of Reward Processing in Addiction : An Image-Based Meta-analysis of Functional Magnetic Resonance Imaging Studies. JAMA Psychiatry, 74(4), 387–398. doi: 10.1001/jamapsychiatry.2016.3084

Mahler, S. V., Brodnik, Z. D., Cox, B. M., Buchta, W. C., Bentzley, B. S., Quintanilla, J., Cope, Z. A., Lin, E. C., Riedy, M. D., Scofield, M. D., Messinger, J., Ruiz, C. M., Riegel, A. C., España, R. A., & Aston-Jones, G. (2019). Chemogenetic Manipulations of Ventral Tegmental Area Dopamine Neurons Reveal Multifaceted Roles in Cocaine Abuse. The Journal of Neuroscience, 39(3), 503–518. doi: 10.1523/JNEUROSCI.0537-18.2018

Michael, D. J., & Wightman, R. M. (1999). Electrochemical monitoring of biogenic amine neurotransmission in real time. Journal of Pharmaceutical and Biomedical Analysis, 19(1-2), 33–46. doi: 10.1016/s0731-7085(98)00145-9

Mittleman, G., Call, S. B., Cockroft, J. L., Goldowitz, D., Matthews, D. B., & Blaha, C. D. (2011). Dopamine dynamics associated with, and resulting from, schedule-induced alcohol self-administration: analyses in dopamine transporter knockout mice. Alcohol, 45(4), 325–339. doi: 10.1016/j.alcohol.2010.12.006

Ostlund, S. B., Wassum, K. M., Murphy, N. P., Balleine, B. W., & Maidment, N. T. (2011). Extracellular dopamine levels in striatal subregions track shifts in motivation and response cost during instrumental conditioning. The Journal of Neuroscience, 31(1), 200–207. doi: 10.1523/JNEUROSCI.4759-10.201

Prater, W., Swamy, M., Beane, M., & Lester, D. (2018) Examining the Effects of Common Laboratory Methods on the Sensitivity of Carbon Fiber Electrodes in Amperometric Recordings of Dopamine. Journal of Behavioral and Brain Science, 8, 117–125. doi: 10.4236/jbbs.2018.83007

Prus, A. J., James, J. R., & Rosecrans, J. A. (2009). Conditioned Place Preference. In J. J. Buccafusco (Ed.), Methods of Behavior Analysis in Neuroscience. (2nd ed.). CRC Press/Taylor & Francis

Robinson, T. E., Gorny, G., Savage, V. R., & Kolb, B. (2002). Widespread but regionally specific effects of experimenter-versus self-administered morphine on dendritic spines in the nucleus accumbens, hippocampus, and neocortex of adult rats. Synapse, 46(4), 271–279. doi: 10.1002/syn.10146

Robinson, D. L., Zitzman, D. L., Smith, K. J., & Spear, L. P. (2011). Fast dopamine release events in the nucleus accumbens of early adolescent rats. Neuroscience, 176, 296–307. doi: 10.1016/j.neuroscience.2010.12.016

Sampedro-Piquero, P., Ávila-Gámiz, F., Moreno Fernández, R. D., Castilla-Ortega, E., & Santín, L. J. (2019). The presence of a social stimulus reduces cocaine-seeking in a place preference conditioning paradigm. Journal of Psychopharmacology, 33(12), 1501–1511. doi: 10.1177/0269881119874414

Schultz, W., Apicella, P., & Ljungberg, T. (1993). Responses of monkey dopamine neurons to reward and conditioned stimuli during successive steps of learning a delayed response task. The Journal of Neuroscience, 13, 900–913. doi: 10.1523/jneurosci.13-03-00900.1993

Schultz, W., Dayan, P., & Montague, P. R. (1997). A Neural Substrate of Prediction and Reward. Science, 275, 1593–1599. doi: 10.1126/science.275.5306.1593

Serafini, R. A., Pryce, K. D., & Zachariou, V. (2020). The Mesolimbic Dopamine System in Chronic Pain and Associated Affective Comorbidities. Biological Psychiatry, 87(1), 64–73. doi: 10.1016/j.biopsych.2019.10.018

Steensland, P., Fredriksson, I., Holst, S., Feltmann, K., Franck, J., Schilström, B., & Carlsson, A. (2012). The monoamine stabilizer (−)-OSU6162 attenuates voluntary ethanol intake and ethanol-induced dopamine output in nucleus accumbens. Biological Psychiatry, 72(10), 823–831. doi: 10.1016/j.biopsych.2012.06.018

Thiel, K. J., Okun, A. C., & Neisewander, J. L. (2008). Social reward-conditioned place preference: A model revealing an interaction between cocaine and social context rewards in rats. Drug and Alcohol Dependence, 96(3), 202–212. doi: 10.1016/j.drugalcdep.2008.02.013

Torquet, N., Marti, F., Campart, C., Tolu, S., Nguyen, C., Oberto, V., Benallaoua, M., Naudé, J., Didienne, S., Debray, N., Jezequel, S., Le Gouestre, L., Hannesse, B., Mariani, J., Mourot, A., & Faure, P. (2018). Social interactions impact on the dopaminergic system and drive individuality. Nature Communications, 9(1), 3081. doi: 10.1038/s41467-018-05526-5

Tye, K. M., Mirzabekov, J. J., Warden, M. R., Ferenczi, E. A., Tsai, H. C., Finkelstein, J., Kim, S. Y., Adhikari, A., Thompson, K. R., Andalman, A. S., Gunaydin, L. A., Witten, I. B., & Deisseroth, K. (2013). Dopamine neurons modulate neural encoding and expression of depression-related behaviour. Nature, 493(7433), 537–541. doi: 10.1038/nature11740

Venniro, M., & Shaham, Y. (2020). An operant social self-administration and choice model in rats. Nature Protocols, 15(4), 1542–1559. doi: 10.1038/s41596-020-0296-6

Venniro, M., Zhang, M., Caprioli, D., Hoots, J. K., Golden, S. A., Heins, C., Morales, M., Epstein, D. H., & Shaham, Y. (2018). Volitional social interaction prevents drug addiction in rat models. Nature Neuroscience, 21(11), 1520–1529. doi: 10.1038/s41593-018-0246-6

Volkow, N. D., & Morales, M. (2015). The Brain on Drugs: From Reward to Addiction. Cell, 162(4), 712–725. doi:10.1016/j.cell.2015.07.046

Yoshizumi, M., Hirao, K., Ueda, K., & Murai, T. (2008). Insight in social behavioral dysfunction in schizophrenia: Preliminary study. Psychiatry and Clinical Neurosciences, 62(6), 669–676. doi: 10.1111/j.1440-1819.2008.01866

